# Encoded cell-material interactions to reroute cytokine signaling for regenerative medicine

**DOI:** 10.1101/2025.10.24.684449

**Authors:** Zachary M. Eidman, Jhanvi Sharma, Joanne C. Lee, Nicholas F. Schulze, Hannah J. Brien, Kevin C. Corn, Marjan Rafat, Jonathan M. Brunger

## Abstract

Regenerative engineering harnesses materials science and stem cell biology to develop strategies to repair damaged and diseased tissue. Despite advances in designer materials, few techniques effectively provide auto-regulated feedback mechanisms that govern how cells sense and respond to discrete microenvironmental changes. Here, we demonstrate that the artificial, juxtacrine-like receptor synthetic Notch (synNotch) can be activated by endogenous multimeric cytokines in solution, without immobilizing materials, revealing a previously unreported activation modality and yielding up to 24-fold dynamic range. To broaden synNotch sensing to monomeric cytokines, we developed nMATRIX, a co-engineered material-cell platform that detects endogenous, soluble ligands and routes them to programmed gene circuits with spatially confined effects. nMATRIX can be tuned to recognize the interleukins IL-1β and IL-6 using synNotch receptors plus cognate biomaterials, yielding up to 68-fold dynamic range and converting these inflammatory inputs into orthogonal outputs that reprogram nearby cell phenotypes. nMATRIX functions across multiple cell types and can incorporate the synNotch-related SNIPR synthetic receptor platform. nMATRIX repurposed inflammatory signals and converted them into anti-inflammatory cues to polarize macrophages (increased CD163, CD206; decreased CD86). Thus, nMATRIX couples native soluble cues to customized cellular responses with tunable sensitivity, offering a flexible materials-based approach for self-regulating regenerative therapies.

## Introduction

Regenerative engineering integrates modern medicine with engineering principles to treat a variety of conditions including injuries, chronic diseases, and age-related degeneration.[1] Recent advances in the areas of biomaterials, stem cell biology and tissue engineering have enabled development of therapies that can repair and regenerate damaged tissue.[2,3] Despite this progress, a number of barriers have prevented these therapies from achieving clinical translation, including immune rejection, imprecise therapeutic control and manufacturing scalability.[4] In chronic inflammatory diseases such as rheumatoid arthritis (RA), immune dysregulation in diseased tissue further complicates the adoption of new therapies.[5,6] Furthermore, current therapeutic platforms lack the capacity to sense and respond to fluctuating biomolecular and mechanical cues, especially in the context of inflammation, which relies on distinct cues to achieve resolution.[7–10] This underscores a pressing need for a new class of auto-regulated and adaptive platforms that can sense dynamic environmental signals and provide context-specific, therapeutic responses to re-establish tissue homeostasis.

Recent developments in materials science have strengthened regenerative engineering efforts by providing new biomaterials that adapt to diverse inputs to coordinate conditional behaviors of cells. These include materials responsive to niche factors such as redox conditions, proteases, temperature, and the biomechanics of the local environment.[11–22] Such materials allow for controlled release of therapeutic agents to elicit favorable cellular responses. Materials have also been developed to adapt based on exogenous inputs, such as light,[23–26] for finely tuned spatiotemporal control over material properties and consequent cell behaviors. While these innovative platforms demonstrate a clear shift toward intelligent therapeutic interfaces, they are restricted to unidirectional modes of engagement; i.e., once remodeled by proteases or ROS-reactivity, or induced to release payloads in response to light or temperature changes, their adaptive features are exhausted, and the materials are unable to continue to instruct cellular behaviors to resolve inflammation or encourage neotissue synthesis.

To overcome limitations associated with material-based strategies, advances in synthetic biology have opened new avenues for engineering cells as therapeutic agents to complement and augment activities enabled via biomaterials that serve as cell delivery vehicles or scaffolds. A diverse repertoire of artificial receptors allows engineered cells to respond to a range of inputs including small molecules, orthogonal proteins, and endogenous cytokines or growth factors.[27–37] Recently, we reported a platform referred to as MATRIX (<u>M</u>aterial <u>A</u>ctivated <u>T</u>o <u>R</u>egulate Inducible gene e<u>X</u>pression),[38] which exploits the synthetic Notch (synNotch) artificial receptor to activate customized transgene expression programs in engineered cells in response to selected inputs presented via functionalized biomaterial surfaces. Similar to native Notch, potent synNotch activation requires ligand immobilization; thus, transgene expression in the MATRIX platform is highly localized and constrained to niches decorated with immobilized activators. Our prior work demonstrated the ability to leverage the MATRIX system to allow engineered cells to respond to soluble, bioinert inputs such as green fluorescent protein (GFP) or mCherry to regulate versatile behaviors relevant to regenerative engineering, including activation of CRISPR-based transcriptome modifiers, modulation of inflammation, and differentiation of human pluripotent stem cells. While the original MATRIX platform was designed to accommodate well-characterized, bioinert ligands, this work expands platform capabilities by converting cytokines such as IL-1 and IL-6 to MATRIX-compatible inputs. Here, we develop a method to reroute actions that cells take in response to these native cytokines. This extension, which we refer to as native MATRIX (nMATRIX), is sensitive to lower input concentrations than orthogonal receptor platforms that have previously been reported and can drive activation levels spanning a wide dynamic range in response to such input. We demonstrate the broad utility of this platform, with high functionality in several cell types relevant to musculoskeletal regenerative medicine, including fibroblasts, mesenchymal stromal cells, macrophages, chondrocytes, and synoviocytes. We further show the ability of this platform to divert the outcome of pro-inflammatory IL-1β signaling, leading such an input to drive the emergence of anti-inflammatory M2-like macrophages rather than the typical, IL-1-generated M1-like pro-inflammatory phenotype. Finally, we demonstrate effective, material-independent synNotch signal transduction for distinct native ligand sub-types via mechanisms that challenge prior reports of synNotch functionality. In doing so, we uncover new rules governing synNotch recognition of native ligands and identify a subset of natural factors that can activate synNotch cells independent of secondary immobilization motifs. Thus, this work informs new strategies for co-engineering cells and biomaterials with synergistic regenerative medicine functionality, providing new insight into receptor design for engineered cell-based therapies.

## Results

### 3.1. Expansion of the MATRIX platform to native factors

To develop a platform that enables potent artificial signaling in response to native growth factors and disease-dysregulated cytokines, we selected a small panel of cytokines including TGF-β1, TNF, IL-1β, and IL-6 as candidates whose signaling could be meaningfully commandeered for regenerative medicine purposes. We derived single chain variable fragments (scFvs) from antibodies previously characterized to bind each of these factors (**Supplementary Table 1**). This panel of scFvs was employed to yield a corresponding suite of receptors to allow engineered MSCs to recognize each cytokine as ligand for synNotch. We first screened whether treatment of any of these MSCs with their respective synNotch ligand resulted in receptor activation in the absence of material surface functionalization. We treated the separate synNotch cells with their respective ligand and measured activity of the synNotch-regulated transgene firefly luciferase. As expected, treatment with soluble IL-1β and IL-6 did not activate corresponding synNotch MSCs. However, we were surprised to find that both TGF-β1 and TNF stimulated artificial signaling in cognate synNotch MSCs to a level of 11-fold and 24-fold, respectively, independent of material-mediated ligand capture and presentation (**Figure 1A**). Anti-VEGF synNotch expressing L929 fibroblasts also demonstrated artificial signaling when supplemented with soluble VEGF in the absence of an immobilizing surface (**Supplemental Figure 1)**. This finding contradicts prior reports describing synNotch as a juxtacrine-like signaling channel that requires ligand immobilization to potentiate signaling and that it is non-responsive to free, soluble inputs. [33,38,39] IL-1β and IL-6 are, like previous inputs we and others have reported on, monomeric signaling factors, while TGFβ-1 TNF, and VEGF are multimeric ligands (dimer, trimer and dimer, respectively[40–42]). Thus, we reasoned that ligand multimerization may enable capture and subsequent presentation to induce synNotch activation independent of biomaterial functionalization. To test this hypothesis, we introduced mutations into enhanced GFP (EGFP), which we previously used for material-dependent signaling via MATRIX, to adapt it into a dimeric ligand (dEGFP).[43] We then treated GFP-sensitive synNotch engineered MSCs with a range of concentrations of dEGFP or wild-type EGFP. The 5 nM concentration of dEGFP rendered dramatic activation of cells expressing the high-affinity, GFP-sensitive LaG16-synNotch (LaG16-Notch) receptor, while wild-type EGFP did not stimulate substantial signaling at any tested concentration (**Figure 1B**), consistent with prior reports [28,38]. We similarly found that dEGFP functions as a productive ligand for KOLF2.1J human induced pluripotent stem cells expressing the LaG16-Notch receptor, suggesting a lack of cell-type dependency on these findings (**Supplemental Figure 2**). These results signify that multimerization converts soluble inputs into those that can productively activate synNotch signaling, while confirming that monomeric inputs such as IL-1β and IL-6 require an alternative mechanism of capture and presentation. We thus pursued IL-1β and IL-6 as native signaling factors compatible with the MATRIX synthetic signaling paradigm.

**Figure 1.**
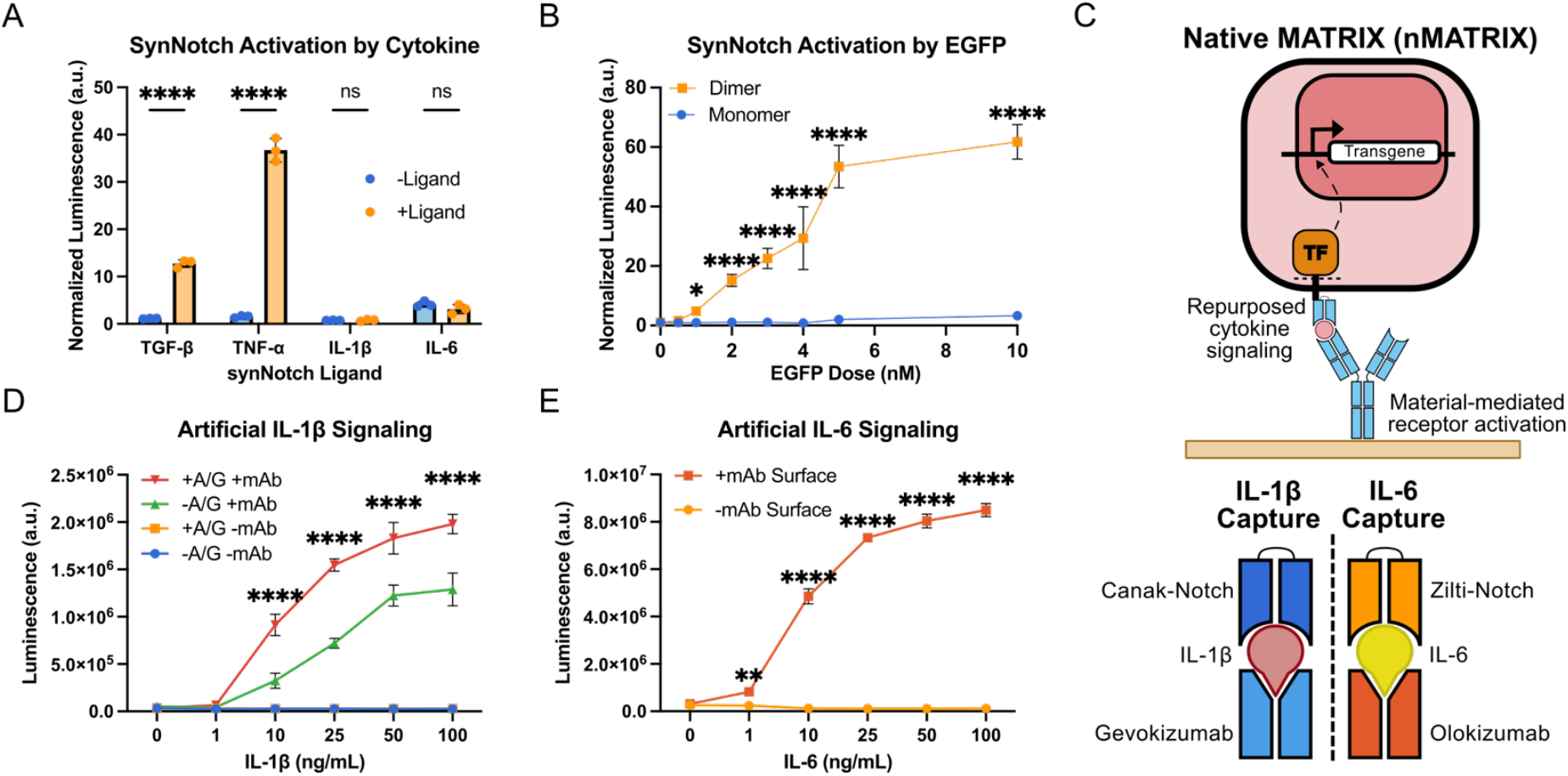
Repurposing MATRIX platform for native inputs. (A) Firefly luminescence normalized to viability indicating synNotch activation in response a panel of synNotch receptors targeted to TGFβ-1, TNF, IL-1β and IL-6. (B) Firefly luminescence normalized to ligand-free conditions indicating Anti-GFP synNotch activation in response to varied doses of monomer versus mutant dimer EGFP. Depicted significance indicated compares +Dimer vs. +Monomer at respective dose. (C) Schematic demonstrating native MATRIX system used to immobilize soluble ligands for presentation to synNotch-engineered cells. (D) Dual binding of either IL-1β or IL-6. Gevokizumab and canakinumab synNotch (Canak-Notch) bind to non-overlapping epitopes of IL-1β. Olokizumab and ziltivekimab synNotch (Zilti-Notch) bind to non-overlapping epitopes of IL-6. (E) Effect of Protein A/G coating on activation of IL-1β-responsive synNotch. Firefly luminescence readout in response to varied doses of IL-1β indicating Canak-Notch activation. Depicted significance indicates +Protein A/G +mAb group compared to each other group at each respective dose (F) Dose response of IL-6-responsive synNotch when plated on olokizumab functionalized surface. Firefly luminescence readout in response to varied doses of IL-6 indicating Zilti-Notch activation. Depicted significance compares -mAb Surface to +mAb Surface for each respective dose. All statistical tests were done using a two-way ANOVA with Tukey’s multiple comparisons. n=3 for all experimental groups *p<0.05, **p<0.01, ****p<0.0001.

To develop a native MATRIX (nMATRIX) platform that can convert IL-1β or IL-6 to inputs for customized, orthogonal signaling, we identified reciprocal binding motifs that could anchor each cytokine via epitopes that do not overlap those recognized by a cognate synNotch receptor (**Figure 1C**). For IL-1β recognition, we again used IL-1-sensitive synNotch cells, engineered to express Canak-Notch, a synNotch receptor utilizing an scFv derived from canakinumab (Canak), a monoclonal antibody which binds IL-1β with high affinity (*K*_d_ = 4.1 nM) and is approved for clinical indications such as systemic juvenile idiopathic arthritis, periodic fever syndrome, and gout flares. To develop a functionalized biomaterial surface that can capture and present IL-1β ligand to Canak-Notch cells, we identified gevokizumab (Gevo), an investigational IL-1β inhibitor (*K*_d_ = 0.29 nM), as a candidate motif, as Canak and Gevo had been previously reported to each recognize distinct, 3D discontinuous epitopes of IL-1β.[44] Thus, to commandeer IL-1β signaling via nMATRIX, we prepared Gevo-functionalized surfaces by either passively adsorbing Gevo directly to cell culture wells or by first coating culture surfaces with protein A/G to orient the Fragment antigen-binding region (Fab) of Gevo away from the culture surface.[45] We then treated cells with IL-1β concentrations ranging from 1-100 ng/ml. While cells on control, Gevo-free surfaces exhibited no Canak-Notch activity when treated with up to 100 ng/ml IL-1β, we observed potent, dose-dependent Canak-Notch activation when cells were cultured on surfaces functionalized with Gevo (**Figure 1D**). At doses of IL-1β frequently used in *in vitro* inflammatory studies (10-100 ng/ml),[46–49] we show responses ranging between 32-fold and 68-fold increase in activity above baseline. As expected, protein A/G-treated surfaces resulted in enhanced Canak-Notch activation as compared to surfaces functionalized via passive adsorption of Gevo, with fold changes in the passive adsorption group peaking at 51-fold change at the 100 ng/ml IL-1β concentration. Thus, by co-engineering Canak-Notch MSCs and a cognate, IL-1-capturing surface, we were able to apply the nMATRIX paradigm to enable material-mediated synthetic signaling in response to a pervasive, pro-inflammatory cytokine.

We similarly developed an nMATRIX variant designed to rewire cellular responses to IL-6. We engineered an IL-6-specific receptor, Zilti-Notch, which borrows variable light and heavy chains from ziltivekimab (Zilti), which blocks IL-6 and IL-6R classical signaling and is thought to bind to the so-called Site I of IL-6 (K_d_=0.52nM). We then selected olokizumab (Olok), which binds to Site III of IL-6 (K_d_=10pM), as the surface-functionalization motif for IL-6 anchorage and display to Zilti-Notch cells.[50,51] AlphaFold-Multimer modeling of IL-6, ziltivekimab scFv, and Olok heavy and light chains also predicted the binding of IL-6 at these sites (**Supplemental Figure 3**). Building off our success with IL-1β recognition, we prepared a protein A/G-coated surface decorated with Olok (**Figure 1C**). We again observed robust dose- and material-dependent responses to a native cytokine via the nMATRIX platform, with Zilti-Notch cells exerting a significant response to levels of IL-6 across the full range of concentrations tested (1-100 ng/ml) (**Figure 1E**). At the 10-25 ng/ml concentration range commonly used for *in vitro* studies that mimic IL-6-driven inflammation, [52–54] Zilti-Notch cells respond with 40-fold to 63-fold above background transgene expression. Altogether, these results suggest that the nMATRIX platform can be generalized across monomeric, soluble inputs for which cognate receptor/material recognition motifs can be identified.

### 3.2. Influence of nMATRIX on cognate cytokine bioactivity

Because both the capture motif and the recognition domain of our synNotch receptors consist of neutralizing antibody fragments, we sought to determine the extent to which the nMATRIX platform may attenuate bioactivity of targeted ligands independent of transgene payload production. Thus, we sought to determine how cytokine retention and bioactivity differ in standard cell culture conditions as compared to cell culture conditions modified with nMATRIX constituents (e.g., functionalized surfaces and synNotch-programmed cells). To do this, we separately treated wild-type (WT) MSCs or Canak-Notch MSCs cultured on either control or Gevo-functionalized surfaces with 10 ng/ml IL-1β. Forty-eight hours later, we collected the culture medium and measured the concentration of IL-1β remaining in the supernatant. Unsurprisingly, we found that Gevo-functionalization dramatically influenced the amount of IL-1β recovered from a well, with IL-1β being undetectable in all supernatants derived from cultures with Gevo-functionalized surfaces (**Figure 2A**). We also found that the level of IL-1β detected in the supernatant of Canak-Notch cells was significantly lower than IL-1β detected in WT, Gevo-free conditions, suggesting that Canak-Notch cells have an enhanced capacity to capture IL-1β as compared to control cells. The nMATRIX variant programmed for IL-6 detection performed similarly, where surface functionalization with IL-6-capturing Olok dominated the bioavailability of IL-6 in supernatant, and Zilti-Notch expression itself also attenuated the amount of IL-6 recovered in the supernatant as compared to control MSCs (**Figure 2B**). Based on these findings, we investigated the degree to which the IL-1β-specific nMATRIX interferes with endogenous cytokine signaling. We programmed a separate line of primary MSCs with an NF-κB transcriptional reporter that regulates expression of a firefly luciferase. We cultured WT or Canak-Notch MSCs plated on control or Gevo-coated surfaces for 24 hours in 0, 1, and 10 ng/ml IL-1β. Cell culture supernatants were then transferred to NF-κB reporter MSCs for five hours before the cells were lysed for a luminescence assay (**Figure 2C**). A subtle but significant reduction in NF-κB transcriptional activity was observed in recipient cells when culture media were derived from Canak-Notch versus WT MSCs (**Figure 2D**). Additionally, the monoclonal antibody surface was unable to completely attenuate NF-κB signaling in the 10 ng/mL group, indicating native signaling can persist at sufficiently high levels. These results suggest that, while Canak-Notch cells effectively bind IL-1β in culture medium, residual cytokine remains highly bioactive. However, consistent with the bioavailability assay, surface Gevo-functionalization dramatically reduced NF-κB transcriptional activity in reporter cells treated with IL-1β conditioned media, indicating that Gevo as employed in these experiments effectively attenuates concentrations of at least 1 ng/ml of IL-1β. Altogether, these results demonstrate that the nMATRIX platform elements can mitigate inflammatory signals in the absence of synNotch-driven, anti-inflammatory transgene expression.

**Figure 2.**
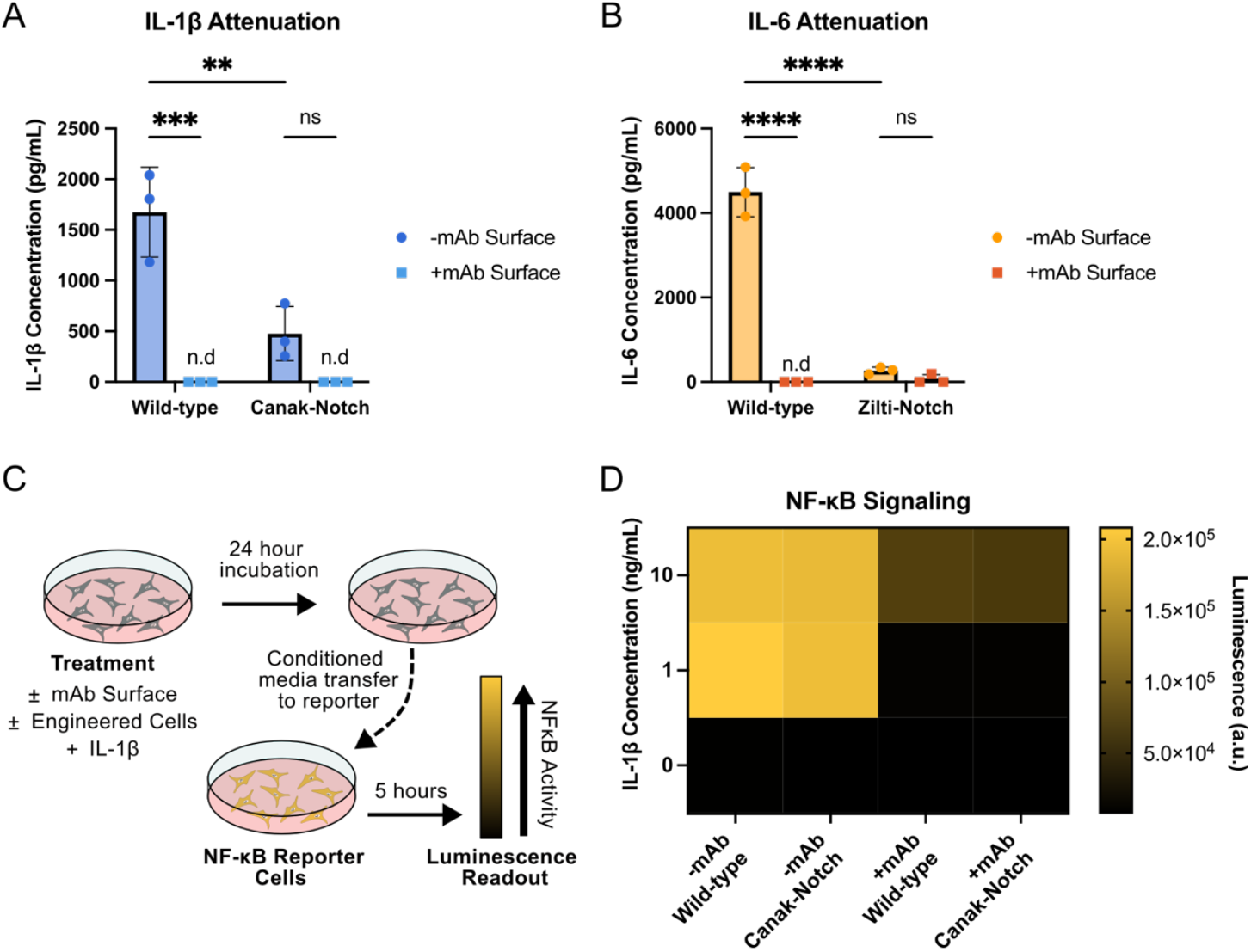
nMATRIX modulates IL-1β and IL-6 levels via receptor and surface neutralization. (A) IL-1β as measured by ELISA 48 hours after supplementing 10 ng/mL to either wild-type mMSCs or Canak-Notch mMSCs with or without the gevokizumab surface. (B) IL-6 as measured by ELISA 48 hours after supplementing 10 ng/mL to either wild-type or Zilti-Notch mMSCs with or without the olokizumab surface. (C) Schematic showing conditioned media treatment of NF-κB reporter cells. (D) Resultant NF-κB signaling 5 hours after transferring IL-1β conditioned media to NF-κB reporter cells. All statistics were performed using a two-way ANOVA with Tukey’s multiple comparisons. n=3 for all experimental groups. **p<0.01 ***p<0.001 ****p<0.0001.

### 3.3 nMATRIX reroutes inflammatory signaling with orthogonal outputs in diverse cell types

We next tested whether the nMATRIX platform is operational in diverse cell types relevant to regenerative engineering. Osteoarthritis, in which inflammatory factors such as IL-1β and IL-6 are known to be dysregulated, represents an area of intense study due to the absence of a disease-modifying therapy. We thus engineered a panel of cells relevant to the homeostasis of articular joints to express the Canak-Notch receptor which regulated expression of firefly luciferase, and assessed their responsiveness to IL-1β with and without Gevo-functionalized surfaces. Previous studies have shown that chondrocytes not only become dysregulated when exposed to proinflammatory cytokines, but they also play an active role in the degradation of cartilage by secreting inflammatory factors.[75] As chondrocytes are also responsible for the production of the cartilage extracellular matrix (ECM), they are an appealing therapeutic target. Indeed, Canak-Notch chondrocytes exhibited activation with as little as 1 ng/mL of IL-1β, with more potent activation seen at higher doses and maximal activation of ∼11-fold over background (**Figure 3A**). Another pathogenic cell type found in the synovium of arthritic joints includes fibroblast-like synoviocytes (FLS), which serve as a source of inflammatory cytokines and proteolytic enzymes attributed to joint ECM degradation.[76] Canak-Notch primary FLS cells showed robust, material-dependent, synNotch activation at doses of 10 ng/mL and 100 ng/mL (**Figure 3B**). Canak-Notch fibroblasts exhibited similar responsiveness to IL-1β, demonstrating significant synNotch activation compared to those plated in either Gevo-free or IL-1-free conditions (**Figure 3C**). Lastly, while infiltrating monocyte-derived macrophages are a hallmark of osteoarthritis and rheumatoid arthritis, prior studies have demonstrated that resident macrophages are a critical modulator in resolving inflammation in affected joints.[55–57] As such, these tissue macrophages are a prime target for engineering to provide more potent therapeutic responses. As a model for these, we used RAW264.7 cells, an immortalized mouse macrophage cell line. Interestingly, though engineered Canak-Notch macrophages exhibited a robust, 66-fold response at the 100 ng/ml IL-1β concentration, the level of induction upon treatment with 10 ng/ml IL-1β was modest relative to responses elicited in MSCs, chondrocytes, fibroblasts, and FLS cells (**Figure 3D**), which may be a result of competition caused by a relatively high expression of the native IL1R1 receptor for IL-1β in these cells.[58] Altogether, these results indicate that cells known to be both sources of and recipients influenced by IL-1β can be successfully programmed via the nMATRIX signaling platform for responding to this pro-inflammatory, arthritis-driving agent.

**Figure 3.**
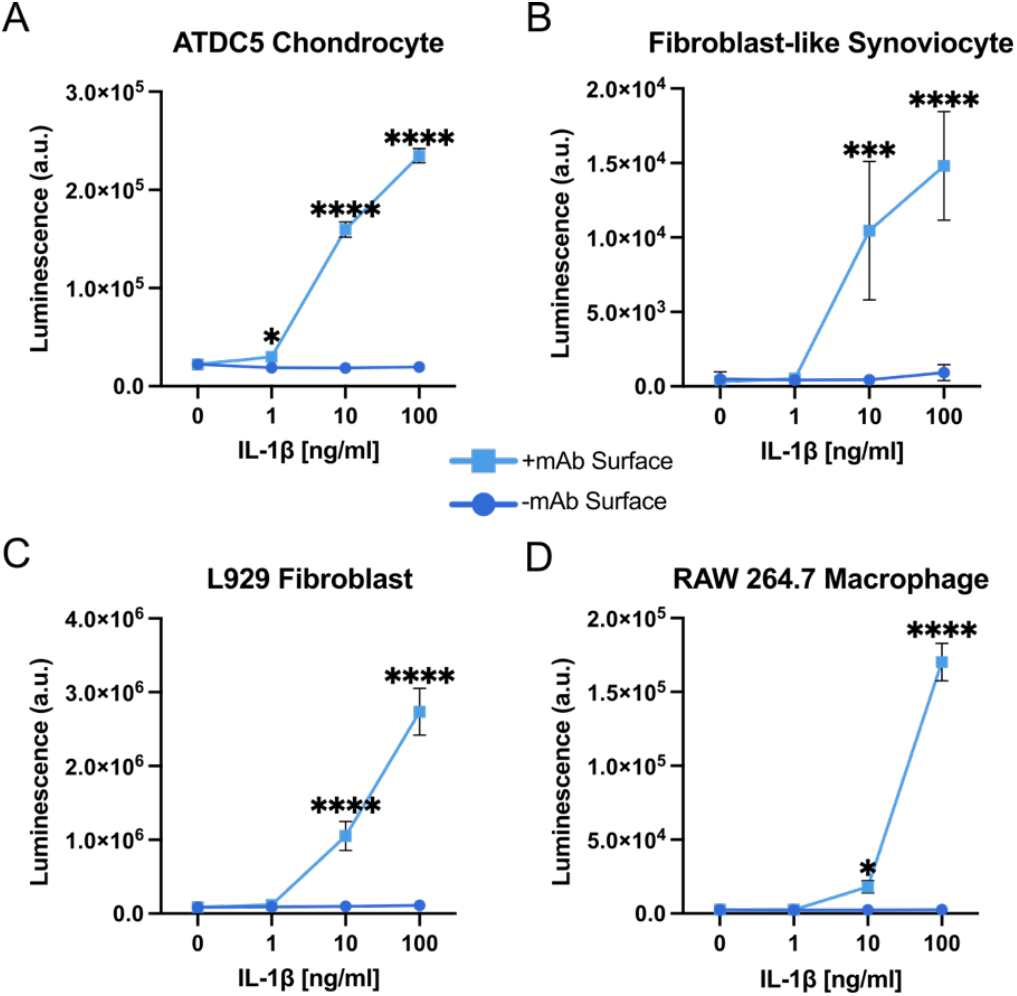
nMATRIX is applicable to different cell types. (A) IL-1β dose dependent activation of (A) ATDC5 chondrocyte-like cells, (B) primary fibroblast-like synoviocyte cells, (C) L929 fibroblasts, and (D) RAW264.7 macrophages. All statistics were performed using a two-way ANOVA with Tukey’s multiple comparisons. n=3 for all experimental groups. *p<0.05, ***p<0.001 ****p<0.0001.

### 3.4 Alternative receptors chassis for nMATRIX signal transduction

The SNIPR platform is an alternative to the classical synNotch regulated intramembrane proteolysis (RIP) receptor.[34] Coding sequences for SNIPRs are smaller than synNotch receptors, making the SNIPR architecture preferred in the context of adeno-associated virus (AAV) gene delivery vectors. Additionally, recent work illustrated that synNotch and SNIPR activation levels differ in the context of soluble, multimeric inputs: SNIPR-engineered T cells activated dramatic transgene expression in response to multimeric inputs such as TGF-β1 due to an endocytic, pH-dependent mechanism that was not active in synNotch T cells.[33] These factors motivated an assessment of whether nMATRIX is compatible with the SNIPR architecture. Thus, we outfitted MSCs with either Canak-SNIPR or Zilti-SNIPR. Results reveal that, despite differences previously reported for SNIPR and synNotch activation mechanisms, the nMATRIX framework accommodates both receptor chassis (**Figure 4A-B**). As with synNotch cells, SNIPR variants demonstrated dependency on both ligand and surface-functionalization for induction. While Canak-Notch activated more potently than Canak-SNIPR on the Gevo-treated surface (**Figure 4A**), Zilti-SNIPR activated more potently than Zilti-Notch when compared to their respective ligand-free and surface-free groups (**Figure 4B**), though both receptors had similar maximal activation levels. Thus, while validating the broad utility of the nMATRIX framework for repurposing IL-1β as an input for programmable artificial signaling, these results also suggest that empirical evaluation may be required to determine whether SNIPR or synNotch should be implemented within the nMATRIX paradigm for each selected ligand.

**Figure 4.**
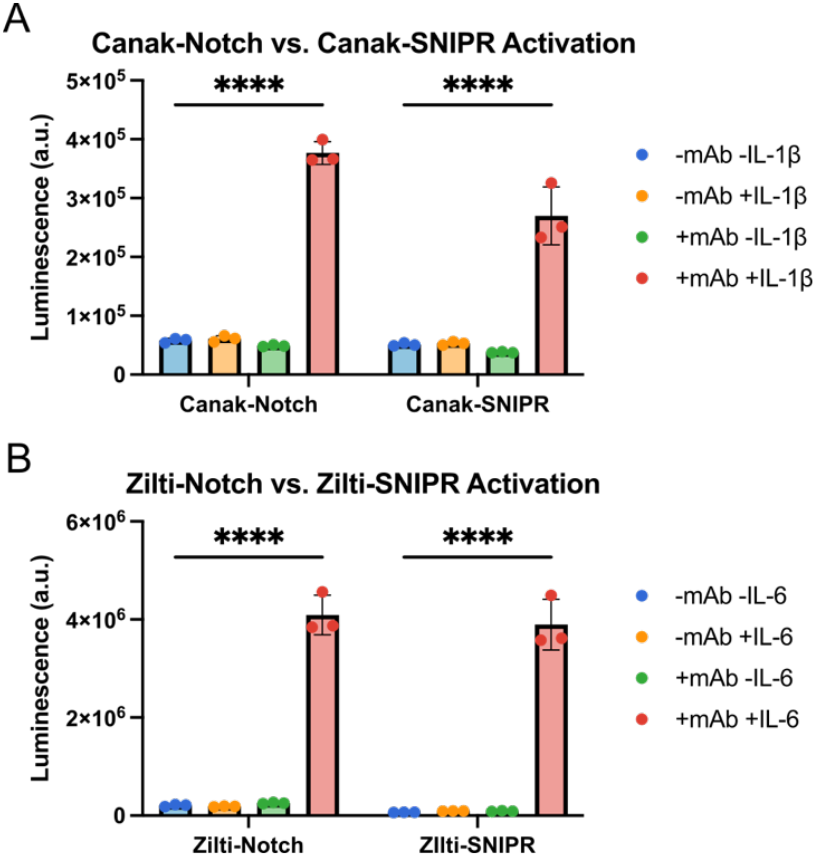
nMATRIX can be adapted to a different receptor platform. (A) SNIPR engineered with canakinumab scFv can be activated in an nMATRIX-dependent manner. (B) SNIPR engineered with ziltivekimab scFv can be activated in an nMATRIX-dependent manner. All statistics were performed using a two-way ANOVA with Tukey’s multiple comparisons. n=3 for all experimental groups. ****p<0.0001.

### 3.5 Synthetic regulation of macrophage polarity via native MATRIX

Having established the ability to commandeer native cytokines for artificial signaling in diverse cell types, we next turned to evaluate whether the nMATRIX platform can effectively redirect cellular responses to pro-inflammatory factors to instead execute anti-inflammatory functions. Due to their association with the pathogenesis and perpetuation of various chronic diseases, we focused on the polarization of macrophages. Whereas M1-like macrophages are considered pro-inflammatory and contribute to autoimmune diseases and inflammatory disorders, M2-like macrophages are considered pro-resolution and aid in restoration of tissue function after injury. A sustained presence of M1-like macrophages, without a transition to M2-like phenotypes, underlies the progression of many chronic degenerative conditions. Thus, we sought to determine whether material-mediated, synNotch-inducible expression of polarizing factors could evoke emergence of an M2-like population. To disentangle the influence of pro-inflammatory cytokines on macrophage polarization, we first returned to our orthogonal MATRIX platform, which relies on bioinert signaling factors such as GFP for induction, to determine whether LaG16-Notch MSCs could induce M2-like polarization of RAW264.7 cells. Thus, we engineered two LaG16-Notch MSC lines: one engineered for inducible expression of control reporter genes mCherry and firefly luciferase (Reporter) and another engineered for inducible expression of the M2-polarizing factors IL-4 and IL-10 (IL4/IL10). We then plated these GFP-responsive MSCs onto control surfaces or surfaces functionalized with the GFP-capturing motif GFP-Trap. Conditioned media were collected and added to RAW264.7 cells at 24- and 48-hour timepoints, and the production of IL-4 and IL-10 was measured 48 hours later via ELISA. While LaG16-Notch IL-4/10 MSCs produced minimal levels of IL-4 or IL-10 when left untreated, we observed >300-fold enhancement in both IL-4 and IL-10 levels in the LaG16-Notch IL4/IL10 group when treating with 5 nM GFP (**Figure 5B,C**). Flow cytometry quantification of M1 and M2 surface markers in macrophages revealed that CD206 was significantly elevated in response to GFP-induced synNotch expression of IL-4 and IL-10 (**Figure 5D, Supplemental Figure 5**). Interestingly, CD163 levels were not upregulated in RAW264.7 cells treated with conditioned medium from GFP-treated LaG16-Notch IL4/10 MSCs (**Figure 5E)**. Congruent with this, prior reports indicated that inflammatory factors are required to elevate expression of CD163 via IL-10 signaling.[59–62] Lastly, MSC culture conditions had little impact on the levels of the M1 marker CD86 (**Figure 5F**) expressed by macrophages. These results demonstrate that synNotch MSC activity can potently influence macrophage phenotype in a ligand-, surface-, and transgene-dependent manner.

**Figure 5.**
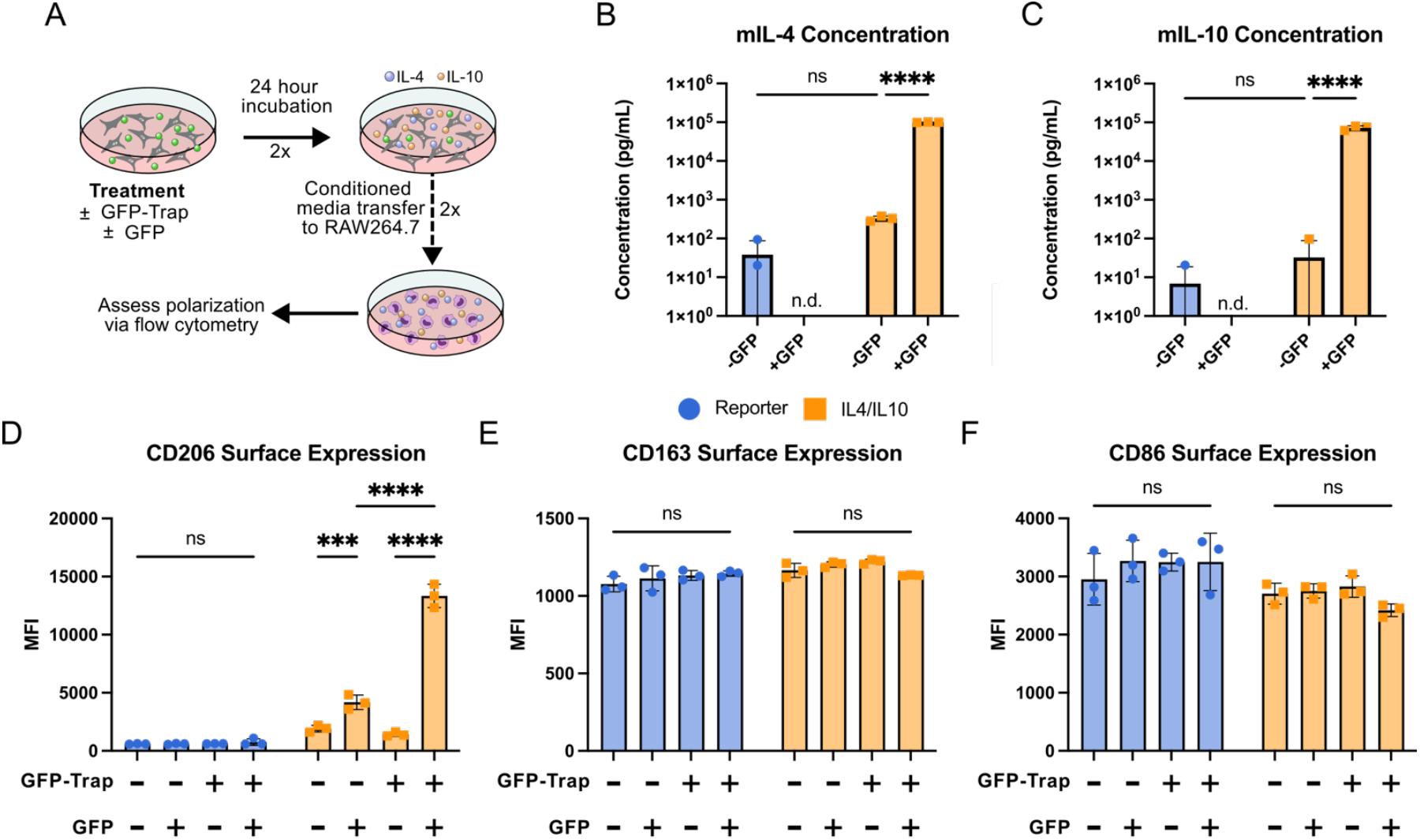
MATRIX can enhance CD206 expression in macrophages (A) Schematic of experimental setup. GFP-responsive synNotch mMSCs containing an inducible reporter transgene or an inducible IL4/IL10 cassette were stimulated with GFP. Conditioned media from these cells was added to RAW 264.7 cells at 24 and 48 hour timepoints. At 72 hours, RAW264.7 surface marker expression was measured via flow cytometry to evaluate polarization. (B-C) Mouse IL-4(B) and mouse IL-10(C) concentrations as measured by ELISA after 24 hours of GFP stimulation of anti-GFP reporter and IL4/IL10 mMSCs. (D-F) Quantified mean fluorescence intensity of CD206(D), CD163(E), and CD86(F) surface expression of RAW264.7 cells after treatment with GFP-induced reporter and IL4/IL10 cell conditioned media. All statistics were performed using a two-way ANOVA (B,C) or three-way ANOVA (D-F) with Tukey’s multiple comparisons n=3 for all experimental groups. ***p<0.001 ****p<0.0001.

Encouraged by these results, we next interrogated whether the nMATRIX platform could reroute IL-1 signaling, which would typically drive a CD86_hi_/CD206_lo_/CD163_lo_ expression pattern, to instead render the inverse phenotype of CD86_lo_/CD206_hi_/CD163_hi_ in macrophages. In this experimental configuration, we employed a transwell co-culture, where macrophages were cultured in the apical chamber and Reporter or IL4/IL10 Canak-Notch-MSCs were cultured in the basal chamber (**Figure 6A**). We stimulated cells with 10 ng/ml IL-β and again observed marked ligand- and surface-dependent upregulation of IL-4 and IL-10 proteins in media collected at Day 3 from Canak-Notch MSCs (**Figure 6B,C**). Flow cytometric results revealed that macrophage phenotype was dramatically influenced by each of surface functionalization, IL-1β treatment, and Canak-Notch MSC transgene payload (**Figure 6D-F, Supplemental Figure 6**). Regardless of the presence of the IL-1 neutralizing Canak-Notch receptor and surface-functionalized Gevo motifs, macrophage CD86 levels were elevated after IL-1β treatment in the presence of Reporter Canak-Notch MSCs. Further, CD206 expression was absent and CD163 levels remained low in co-cultures with Reporter Canak-Notch MSCs. In contrast, IL-1β treatment in co-cultures with IL4/IL10 Canak-Notch MSCs resulted in slight but significant elevation of CD86 only in the synNotch-off, Gevo-free state. When Gevo-functionalization enables ligand-dependent production of IL-4 and IL-10 by Canak-Notch MSCs, CD86 levels are completely attenuated. As in the prior GFP MATRIX experiment, we observed elevated levels of CD206 in response to synNotch-driven IL-4 and IL-10 production, despite the presence of pro-inflammatory IL-1β. Furthermore, in the presence of IL-1β and resultant payload transgenes from IL4/10 Canak-Notch MSCs, we observed dramatically enhanced CD163 expression. Taken together, these results illustrate that the nMATRIX platform with Canak-Notch MSCs reroutes the M1-like outcome of pro-inflammatory IL-1β signaling to drive macrophage polarization.

**Figure 6.**
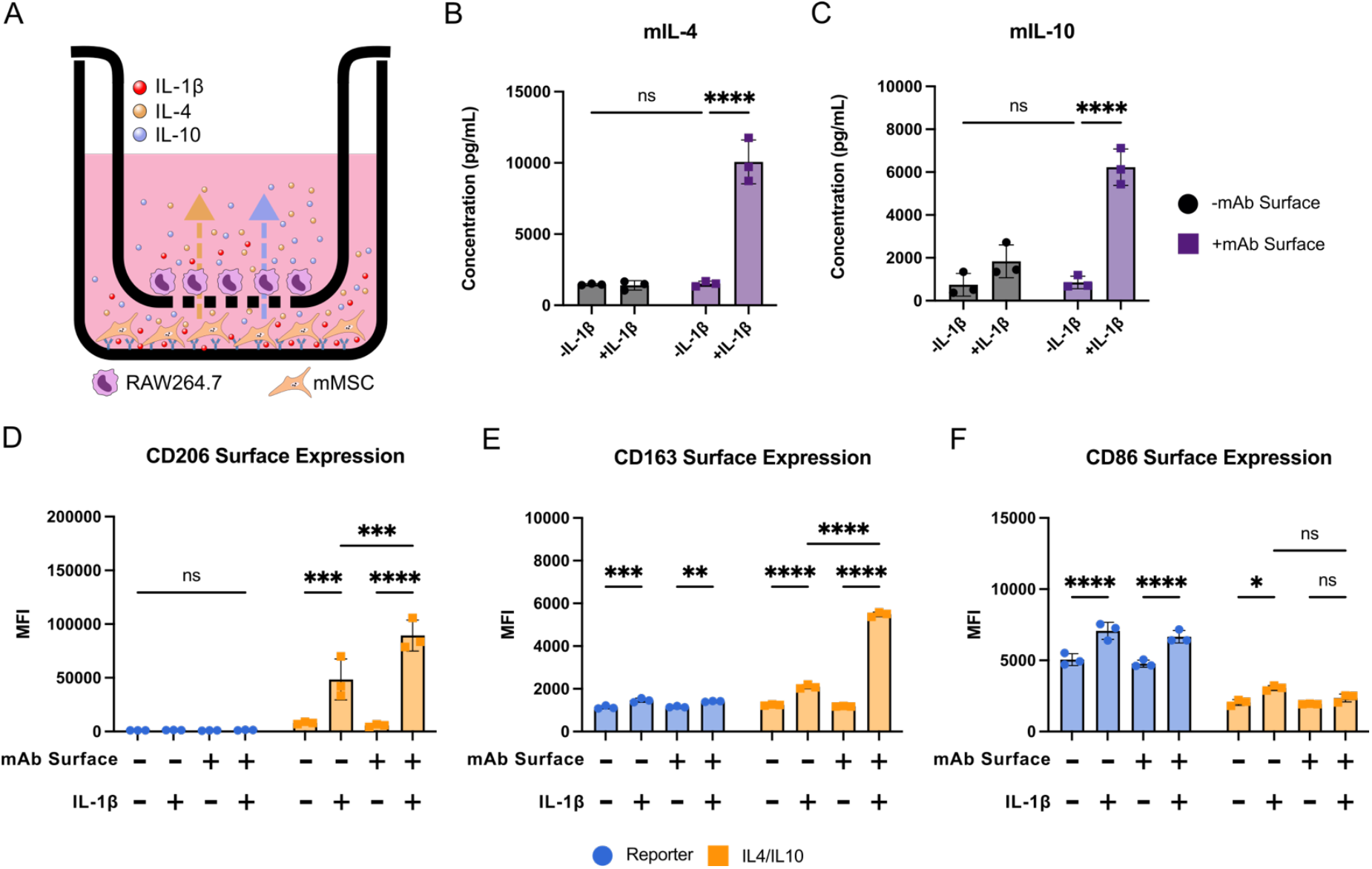
nMATRIX can polarize macrophages toward M2-like phenotypes. (A) Schematic of transwell setup. Canak-Notch synNotch mMSCs containing an inducible reporter transgene or an inducible IL4/IL10 cassette were stimulated with 10 ng/mL IL-1β. After 24 hours of mMSC stimulation, RAW264.7 cells in the apical transwell chamber were transferred into the basal mMSC chamber. After 48 hours of co-culture, RAW264.7 surface marker expression was measured via flow cytometry to evaluate polarization. (B-C) Murine IL-4 (B) and murine IL-10 (C) concentrations as measured by ELISA after co-culture with RAW264.7 cells. (D-F) Quantified mean fluorescence intensity of CD206 (D), CD163 (E), and CD86 (F) surface expression of RAW264.7 cells after treatment with IL-1β-induced reporter and IL4/IL10 co-culture. All statistics were performed using a two-way ANOVA (B,C) or three-way ANOVA (D-F) with Tukey’s multiple comparisons. n=3 for all experimental groups. *p<0.05 **p<0.01 ***p<0.001 ****p<0.0001.

## Discussion

Here, we demonstrate that our previously described MATRIX platform, comprised of designer biomaterial surfaces and engineered cells, can be configured to sense and respond to native, soluble molecules. Notably, nMATRIX utilizes customizable, orthogonal synNotch receptors that induce expression of downstream gene circuits, allowing for the conversion of various pro-inflammatory cytokines into alternative outputs, without influencing endogenous pathways. Our platform enables robust, inducible gene expression with high fold-change activation within a biologically relevant range of cytokine concentrations.[46–49,52–54] Both the material and receptor components of nMATRIX have demonstrated cytokine neutralizing capabilities, which can be used to modulate endogenous inflammatory signaling. Additionally, nMATRIX acts as a Boolean AND gate, requiring engineered cells, ligand and the immobilizing surface to be present to produce programmed outputs, allowing for spatiotemporally controlled therapeutic delivery. nMATRIX offers broad applicability, as it can be used to produce artificial signaling in a wide variety of cell types implicated in musculoskeletal diseases including mesenchymal stem cells, macrophages and fibroblast-like synoviocytes, and can be used with alternative synthetic receptor architectures such as SNIPR. Lastly, we demonstrated nMATRIX can be used to convert the highly inflammatory signal IL-1β into an anti-inflammatory output to polarize macrophages towards a more anti-inflammatory phenotype. Overall, nMATRIX is a highly customizable platform that allows for material-dependent, prescribed cellular programming in response to elevated levels of native factors that contribute to inflammatory diseases.

As the immune system plays a significant role in the progression and chronic dysregulation of inflammatory diseases, environmentally responsive biomaterials, including hydrogels[66–69], polymeric scaffolds[67–69], and nanoparticles[70–73], have been developed to modulate the immune environment by delivering immunomodulatory agents in a regulated manner. Although these technologies initially offer improved control and sustained release of therapeutics, this effect dissipates as the respective material degrades and releases its payload. Engineered cell-based therapies offer a potential solution for auto-regulated and autonomous immunomodulation, as cells can be tuned to effectively sense and respond to distinct cues in the microenvironment. [74–81] Here, we demonstrate that nMATRIX can detect inflammatory cues and respond with anti-inflammatory cues, influencing the phenotype of nearby cells and pushing them towards a more pro-resolution fate. Additionally, both components of nMATRIX have cytokine neutralizing capabilities, enhancing the therapeutic potential of the platform beyond cellular outputs. As such, nMATRIX can potentially be used to resolve dysregulation in diseased tissue by combining inflammatory knockdown and regulated cell feedback to regain tissue homeostasis.

Macrophages are a highly plastic cell type with the ability to change phenotype based on cues in the microenvironment. In rheumatoid arthritis, inflammatory macrophages contribute to both the initial onset and progression of disease through production of cytokines such as TNF, IL-1β and IL-6.[82] As such, repolarization of macrophages has become a focus in potential treatments of numerous diseases from cancer to arthritis.[83] Previous studies have used hydrogels or polymeric scaffolds with inherent physical properties that drive this M2 polarization[84–87]. Similarly, mRNA-, plasmid-, or protein-loaded materials[88–91] have been shown to effectively immunomodulate macrophages by delivering anti-inflammatory cytokines like IL-4 and IL-10. This methodology has also been applied by engineering cells to overexpress anti-inflammatory cytokines like IL-10.[92,93]. Though these studies have shown efficacy in polarizing macrophages towards an anti-inflammatory phenotype, the platforms function unidirectionally and cannot autonomously sense external stimuli to generate customized responses. Here, we demonstrated nMATRIX can effectively reroute IL-1β signaling to instead produce IL-4 and IL-10 via artificial gene circuits in a material- and ligand-dependent manner, which steered macrophages towards a more regenerative, anti-inflammatory phenotype. This provides an increased benefit over the previously described platforms in that nMATRIX is an autoregulated system that responds to dynamic cues in the microenvironment. The modular programmability of nMATRIX suggests the opposite could be implemented as well: pushing an M2-like macrophage towards an M1 phenotype. This would be especially useful in a tumor microenvironment, as various studies have shown polarizing tumor-associated macrophages with inflammatory cytokines can improve immune response to cancer.[94]

A core principle of synthetic biology is developing modular frameworks that can utilize native inputs to drive desired cellular outputs. Early synthetic receptors like chimeric antigen receptors (CARs) fuse elements from the endogenous T cell receptor (TCR) signaling apparatus to synthetic binding domains, enabling activation of T cells in response to selected surface marker exhibition.[95] Although CARs have been shown to effectively activate T cells in response to surface markers such CD19 and HER2[96,97], and even a soluble, multimeric factor, TGF-β[98], their outputs are limited to endogenous TCR signaling pathways. Other approaches including chimeric cytokine receptors (ChR) and synthetic cytokine receptors (SyCyRs) hijack native cytokine receptor signaling pathways by modifying the ectodomain of a cytokine receptor to activate via different cytokine input or a non-physiological input such as GFP. [99,100] These technologies have shown great promise, but their signaling mechanism are susceptible to crosstalk and have limited applicability to other cell types. Another modular receptor system, MESA, has demonstrated artificial, orthogonal signaling utilizing dimerizing ligands such dTomato or VEGF.[29,101] In this work, we have demonstrated synNotch can also be activated using soluble, multimeric cytokines like TNF, TGF-β1 and VEGF. This contradicts previous reports showing that TGF-β could not be used to activate synNotch, indicating a potential cell type dependency for this interaction (e.g. adherent vs. suspension cells).[33] nMATRIX further adds to the synthetic biology toolbox by enabling synNotch signaling with soluble, monomeric, native inputs like IL-1β and IL-6 via biomaterial immobilization. Furthermore, we demonstrated nMATRIX can act as a cytokine converter via orthogonal signaling pathways that can be modulated based on desired application. We also showed that nMATRIX effectively signals with other RIP receptors like SNIPR, suggesting the platform can be used with other juxtacrine receptor formats. Lastly, the dual epitope binding methodology applied in this work could potentially facilitate monomeric signaling in other receptor architectures like MESA and CARs by outfitting them with our cognate scFv pairs.

Although a flexible and effective platform, nMATRIX in its current form has various limitations. As material-mediated ligand immobilization is required for effective signaling, the system would need to be adapted into a 3D model for *in vivo* applications. Antibody conjugated hydrogels have been previously described, and would be a strong candidate for delivering both immobilizing antibody and engineered cells *in vivo*. [102,103] Additionally, the antibodies described in this work are humanized and have limited cross-reactivity with murine cytokines, and preclinical trials would need to use humanized IL-1β or IL-6 mouse models. Lastly, since nMATRIX requires two affinity motifs that bind to distinct epitopes on the same molecule simultaneously, it would likely be difficult to adapt to poorly characterized molecules with few targeting motifs. Despite these limitations, nMATRIX remains a highly flexible, modular platform with broad use for cellular programming. Future experiments could explore porting nMATRIX into 3D hydrogels for *in vivo* applications and developing cell lines that co-express cognate receptors to enable material-free signaling.

In conclusion, we present the nMATRIX platform, a hybrid system integrating material-guided signal transduction with computational capabilities of engineered cells. Crucially, the platform confines both sensing and effector activity to biomaterial-contact regions, minimizing off-site effects. The platform is also highly modular: both the cells and material surfaces can be programmed to recognize and respond to virtually any selected ligand, enabling precise control over cellular behavior. This adaptability is especially valuable in complex pathological environments such as solid tumors, chronic inflammatory conditions, and degenerative tissues, where coordinated, multi-cellular intervention is often required. By harnessing the native capabilities of diverse cell types and augmenting them with nMATRIX, it becomes possible to design multi-layered therapeutic systems capable of sensing and responding to their surroundings with high precision. In summary, our platform provides a robust method of producing auto-regulated and orthogonal signaling in response to physiological inputs like cytokines.

## Materials and Methods

### Plasmids and Cloning

Plasmids were constructed using NEB HiFi DNA Assembly Mix (New England Biolabs) and were subsequently transformed into DH5α competent cells. Cultures were plated on LB Agar + Ampicillin (Sigma) and grown overnight at 37°C. Colonies were picked and validated via whole plasmid sequencing (Plasmidsaurus).

### Virus production

Lentivirus was produced by transfecting Lx293T cells (Clontech) with 2 μg of transfer vector, 1.5 μg of pCMV-dR8.91 packaging vector[104] and 0.6 μg of pMD2.G envelope vector[105] (Addgene #12259) using Lipofectamine 3000 (Thermo Scientific) following manufacturer’s instructions. 16 hours later, medium was exchanged for fresh DMEM-High Glucose supplemented with GlutaMAX and sodium pyruvate + 10% heat-inactivated FBS. 24 and 48 hours later, viral supernatant was collected, filtered with a 0.45 μm PVDF filter (CELLTREAT) and pooled for cell transduction or stored short-term (<1 week) at 4°C or long-term at −80°C.

### Dimeric EGFP (dEGFP) production and purification

#### dEGFP Cloning

A mutated superfolder GFP sequence was used as the basis of the GFP dimer for copper-mediated disulfide-based dimerization, based on a previously reported amino acid sequence [234,235]. Two naturally occurring cysteines, C48 and C70, were mutated to alanine to reduce off-target disulfide bonds, while the aspartic acid residue, D117, was mutated to cysteine for disulfide bonding. In addition, a His-tag was added for protein purification. The mutated superfolder GFP coding sequence was produced through Thermo Fisher Scientific’s GeneArt service and cloned into a pBH4 backbone for protein production.

#### Protein expression

Plasmids containing the mutant GFP dimer sequence were transformed into BL21-DE3 *E. coli* competent cells (NEB) and plated on an LB+agar with ampicillin supplementation plate for overnight incubation at 37°C. Colonies were then picked for three 4 mL starter cultures to be grown with ampicillin supplementation overnight at 37°C. The next day, the starter cultures were added into 1L of LB broth supplemented with ampicillin and allowed to grow at 37°C until the OD was measured in the 0.6-0.8 range. The culture was then induced with 1 mM IPTG (Gold Bio) and incubated for another 4 hours at 37°C. Then the culture was spun down at 4000x*g* for 20 minutes to harvest the cells. The pellets were then consolidated into 5 mL of PBS and then spun down again at 4000x*g* for 10 minutes. Pellets were frozen and stored at -20°C for up to 2 months before purification.Cell pellets were lysed using the BugBuster (Novagen) reagent. Briefly, the wet weight of the pellet was determined, and 5 mL of BugBuster reagent was added per 1.5 g of pellet. 10 µL of Lysonase reagent was then added to the suspension, which was then incubated at room temperature for 30 minutes on a rotating mixer. To remove insoluble cell debris, the suspension was then centrifuged at 4000x*g* for 1 hour at 4°C.

#### Protein purification

His-tagged protein was isolated using a cobalt resin column. After the suspension was passed through the gravity column, the column was washed with one column volume wash buffer (20 mM Tris-HCl, 150 mM NaCl, 10 mM imidazole). The protein was then collected in fractions using elution buffer (20 mM Tris-HCl, 150 mM NaCl, 250 mM imidazole). Fractions were pooled and then concentrated into 1 mL buffer (20 mM Tris pH 9.0, 100 mM NaCl) [235] using Amicon molecular weight cutoff (MWCO) filters (Sigma-Aldrich).

#### Dimerization

The GFP protein was then induced to form disulfide bonds using 10 mL dimerization buffer (20 mM Tris pH 9.0, 100 mM NaCl, 5 mM CuSO4). The reaction was allowed to proceed for 15 minutes at room temperature until quenching by addition of 50 mM EDTA. The protein was then exchanged into anion exchange buffer (10 mM Tris pH 9.5, 1 mM EDTA) using Amicon MWCO filters. Using an anion exchange column (Cytiva), the dimers were purified from the monomers via a salt gradient of 0-1 M NaCl in anion exchange buffer [235]. Fractions were collected and validated using non-reducing SDS-PAGE. Fractions containing GFP dimers were desalted into DPBS using Amicon MWCO filters and aliquoted and frozen at -80°C.

### Cell Culture

#### Murine Mesenchymal Stem Cell

Bone-marrow derived C57BL/6 mMSCs (Cyagen) were cultured in MEM-α + GlutaMAX (Gibco) supplemented with 15% FBS in an incubator at 37°C and 5% carbon dioxide. For routine subcultivation, cells were rinsed with 1X DPBS (Gibco) and treated with TrypLE for five minutes at 37°C to detach cells. After quenching cells with complete medium, they were spun at 300xg for 5 minutes and replated in culture medium.

#### L929 Fibroblasts

Mouse L929 fibroblasts (ATCC# CCL-1) were cultured in DMEM + GlutaMAX (Gibco) supplemented with 10% heat-inactivated FBS in an incubator at 37°C and 5% carbon dioxide. For routine subcultivation, cells were treated with TrypLE for five minutes at 37°C to detach cells. After quenching cells with complete medium, they were spun at 300x*g* for 5 minutes and replated in culture medium.

#### RAW 264.7 Macrophages

RAW 264.7 macrophages (ATCC# TIB-71) were cultured in DMEM + GlutaMAX (Gibco) supplemented with 10% FBS in an incubator at 37°C and 5% carbon dioxide. For routine subcultivation, cells were rinsed with 1X DPBS (Gibco) and treated with TrypLE for five minutes at 37°C to detach cells. After quenching cells with complete medium, they were spun at 300x*g* for 5 minutes and replated in culture medium.

#### ATDC5 Chondrocytes

ATDC5s (Sigma Aldrich) were cultured in DMEM:F12 + GlutaMAX (Gibco) supplemented with 5% FBS in an incubator at 37°C and 5% carbon dioxide. For routine subcultivation, cells were rinsed with 1X DPBS (Gibco) and treated with TrypLE for five minutes at 37°C to detach cells. After quenching cells with complete medium, they were spun at 300x*g* for 5 minutes and replated in culture medium.

#### Fibroblast-like Synoviocytes

Primary fibroblast-like synoviocytes were harvested from 8 week old C57BL/6 mice according to the following protocol.[106] Isolated FLS cells were cultured on adsorbed rat tail collagen I (Gibco) in DMEM + GlutaMAX (Gibco) supplemented with 10% FBS in an incubator at 37°C and 5% carbon dioxide. For routine subcultivation, cells were rinsed with 1X DPBS (Gibco) and treated with TrypLE for five minutes at 37°C to detach cells. After quenching cells with complete medium, they were spun at 300x*g* for 5 minutes and replated on rat tail collagen I in culture medium.

#### Lentiviral Transduction

All cell lines were produced using reverse transduction using virus concentrated with LentiX Concentrator (Takara). Viral media was supplemented with 4 μg/mL of polybrene to enhance transduction efficiency and incubated with cells overnight at 37°C before being replaced with fresh cell culture media. Cells were selected using 10 μg/mL of puromycin (Sigma) when applicable. Unless otherwise noted, all cells following transduction were sorted for synNotch and transgene expression via fluorescence activated cell sorting (FACS).

#### Material-free activation of cytokine responsive synNotch receptors

mMSCs were co-engineered with synNotch receptors with scFvs that each bind to one of TNF-α, TGF-β1, IL-1β or IL-6 and a transgene cassette expressing a tetracycline response element (TRE) driven mCherry and luciferase output. 8,000 cells were plated in a white-walled 96-well plate (Corning) and treated with their respective ligands: TNF-α (25 ng/mL, StemCellTech), TGF-β1 (50ng/mL, StemCellTech), IL-1β (10 ng/mL, MedChemExpress) or IL-6 (10 ng/mL, MedChemExpress). After 48 hours, CellTiter-Fluor (Promega) was performed according to manufacturer’s instructions and read on a Tecan Infinite M1000 Pro plate reader. Following this, a BrightGlo (Promega) assay measuring firefly luminescence as a measure of synNotch activity was performed according to manufacturer’s protocol on a Tecan Infinite M1000 Pro plate reader. Normalization was performed by dividing luminescence values each well’s respective CellTiter-Fluor output.

#### GFP Dimer Activation Assay

synNotch L929 fibroblasts were plated at a density of 18,000 cells/well (56,250 cells/cm2) on a control surface of streptavidin only. The LaG16 (K_d_ of 0.7 nM) synNotch receptor was exposed to a range of concentrations from 0 to 100 nM of either recombinant GFP purified from *E. coli* via immobilized metal affinity chromatography or mutant purified GFP dimer (purification process described above). After 48 hours, firefly luminescence was measured using the BrightGlo luminescence assay (Promega) on a Tecan Infinite M1000 Pro plate reader. Results are expressed as fold change relative to GFP-free conditions.

#### Antibody surface functionalization

Recombinant protein A/G (ThermoFisher) was reconstituted according to manufacturer’s recommendations. Aliquoted Protein A/G was further diluted to 10ug/mL in BupH Carbonate-Bicarbonate buffer (ThermoFisher) and 100ul per well was added to tissue culture (TC) treated 96-well plates (Corning). After incubating 1 hour at room temperature (RT), the solution was aspirated and washed with 200 μl of 1X DPBS (Gibco). Gevokizumab (MedChemExpress) or Olokizumab (MedChemExpress) were diluted to 10 μg/mL and 100 μl were added to the Protein A/G coated wells and left to coat overnight at 4°C. The following day, wells were washed with 200 μl of 1X DPBS and used immediately.

#### Biomaterial based activation of synNotch

After functionalizing a plate with gevokizumab, 12,000 canakinumab synNotch engineered mMSCs were plated in each well and supplemented with media containing IL-1β in a range of concentrations from 0-100 ng/mL. After 48 hours, firefly luminescence was measured via BrightGlo (Promega) luminescence assay on a Tecan Infinite M1000 Pro plate reader. To assess IL-6 synNotch activation, this same procedure was performed using an olokizumab functionalized surface and ziltivekimab engineered synNotch cells dosed with 0-100 ng/mL of IL-6.

#### IL-1β ELISA

96-well TC plates were coated with Protein A/G and gevokizumab as previously mentioned. 12,000 of either wild-type or canakinumab synNotch engineered cells were plated in each well and dosed with 10 ng/mL of IL-1β. After 48 hours of culture, conditioned media were spun at 1500x*g* for 15 minutes and were analyzed using an IL-1β ELISA kit (R&DSystems DY201) according to the manufacturer’s protocol.

#### IL-6 ELISA

96-well TC plates were coated with Protein A/G and olokizumab as previously mentioned. 12,000 of either wild-type or ziltivekimab synNotch engineered cells were plated in each well and dosed with 10 ng/mL of IL-6. After 48 hours of culture, conditioned media was spun at 1500x*g* for 15 minutes and was analyzed using an IL-6 ELISA kit (R&DSystems DY206) according to the manufacturer’s protocol.

#### *NF-κB Transcriptional Activity* Assay

NF-κB reporter mMSCs were generated via lentiviral transduction of a plasmid encoding 4 repeats of the NF-κB response elements followed by a minimal promoter sequence,[107] allowing for activation of a downstream luciferase transgene when NF-κB is active. To assess modulation of native NF-κB signaling, 96-well TC plates were treated with Protein A/G and gevokizumab as previously described and 12,000 of either wild-type or canakinumab engineered reporter mMSCs were added to each well and supplemented with 0, 1, or 10ng/mL of IL-1β in 100 μl of media. After 24 hours, conditioned media from these groups were added to NF-κB reporter mMSCs, which were plated at a density of 8,000 cells per well in a 96-well, white-walled TC treated plate the day prior. After 5 hours of co-culture, a BrightGlo (Promega) luminescence assay was performed on the NF-κB reporter cells to quantify NF-κB activity.

#### SynNotch and SNIPR activation

IL-1β and IL-6 SNIPR receptors (Canak-SNIPR and Zilti-SNIPR, respectively) were created by replacing the transmembrane domain of their synNotch counterparts with the CD8 Hinge, Notch 1TM and Notch2JM domains derived from an anti-CD19 SNIPR construct (Addgene #188375)[34] via PCR and Gibson assembly. A flexible (G3S)x3 linker was added between the scFv and CD8 Hinge domains for additional flexibility. 12,000 mMSCs expressing either canakinumab engineered synNotch or canakinumab engineered SNIPR and a reporter luciferase transgene were plated on a white-walled, 96-well plate. Wells were treated with gevokizumab as previously described. Cells were treated with 10ng/mL of IL-1β and a BrightGlo luminescence assay was performed 48 hours after plating to assess receptor activity. The same procedure was performed to assess IL-6 mediated activity of ziltivekimab engineered synNotch and SNIPR cells plated on olokizumab functionalized surfaces.

#### GFP mediated polarization

anti-GFP LaG16-synNotch engineered cells were transduced with a synNotch inducible transgene circuit that expresses mouse IL-4 and mouse IL-10 in response to activation. Wells of a 24 well non-TC plate were coated with GFP-TRAP-PEG as previously described[38] and 80,000 mMSCs were plated and treated with 5 nM GFP. In a separate TC-treated 24-well plate, 180,000 RAW264.7 macrophages were plated. After 24 hours, conditioned media from the anti-GFP mMSCs were spun down at 1500x*g* for 10 minutes and added to the RAW264.7 cells. Media without GFP were added to the anti-GFP mMSCs and were allowed to incubate another 24 hours before being transferred to the RAW264.7 macrophages as previously described. After an additional 24 hours of incubation, RAW264.7 were lifted using Accutase and labeled for CD86, CD163 and CD206 surface markers using antibodies listed in Supplementary Table 2 before undergoing flow cytometry.

#### IL-1β mediated polarization

Canakinumab-synNotch mMSCs were engineered with the mouse IL-4/IL-10 payload via lentiviral transduction as described earlier. 80,000 mMSCs were added to the basal chamber of a gevokizumab functionalized transwell plate (Corning) and were treated with 10 ng/mL of IL-1β. Simultaneously, 30,000 RAW264.7 macrophages were added to the apical chamber of a transwell and kept in a separate well with 500 μl of media. After 24 hours of incubation, the RAW264.7 transwell inserts were transferred into the mMSC wells and allowed to incubate for an additional 48 hours. After co-culturing, RAW264.7 cells were lifted using Accutase and stained for CD86, CD163 and CD206 surface markers before undergoing flow cytometry.

#### mIL-4 and mIL-10 ELISA

Mouse IL-4 and mouse IL-10 ELISA levels were assessed using respective ELISA kits (R&DSystems, DY404 and DY417) according to manufacturer instructions.

#### Flow Cytometry

All quantitative flow cytometry was performed on Amnis Cellstream flow cytometer. Cells were resuspended in DPBS supplemented with 2% FBS. Fluorescence activated cell sorting (FACS) was performed at the Vanderbilt Flow Core using a BD 5 Laser FACS Aria. Antibodies were diluted according to manufacturer’s instructions for all samples. A list of antibodies used can be found in Supplemental Table 2.

#### Statistical analysis

Graphs were produced using GraphPad Prism 10. All bar graphs display the mean of triplicate values with errors bars showing the standard deviation. To determine significance in experiments, a two- or three-way ANOVA with an alpha of 0.05 was used to compare between groups as needed. ANOVA tests were conducted along with Tukey’s post-hoc test with multiple comparisons.

## Acknowledgements

The authors acknowledge funding support from NSF Career Award CBET-2237639, NIH R01AR083437, and the Vanderbilt Institute for Clinical and Translational Research (VR72804). This work was also supported by the Arthritis National Research Foundation Judy E. Green Valiant Women’s Fellowship. Fluorescence activated cell sorting was performed at the Vanderbilt Flow Cytometry Shared Resource, which is supported by the Vanderbilt Ingram Cancer Center (P30 CA068485) and the Vanderbilt Digestive Disease Research Center (DK058404). We thank Megan Keech and Larry Stokes for technical assistance.

