## Supplemental File for "Encoded cell-material interactions to reroute cytokine signaling for regenerative medicine"

**Supplemental Materials and Methods**

*Material-free activation of VEGF-responsive synNotch receptors*: L929 fibroblasts were co-engineered with synNotch receptors with a VEGF binding scFv (Supplemental Table 1) and a transgene cassette expressing a tetracycline response element (TRE) driven luciferase output. 9,000 cells were plated in a white-walled 96-well plate (Corning) and treated with 25 ng/ml of VEGF (StemCellTech). After 48 hours, CellTiter-Fluor (Promega) was performed according to manufacturer's instructions and read on a Tecan Infinite M1000 Pro plate reader. Following this, a BrightGlo (Promega) assay measuring firefly luminescence as a measure of synNotch activity was performed according to manufacturer's protocol on a Tecan Infinite M1000 Pro plate reader. Normalization was performed by dividing luminescence values each well's respective CellTiter-Fluor output.

*Maintenance of KOLF2.1J induced pluripotent stem cells*: KOLF2.1J[10] stem cells were maintained in StemFlex (Gibco) in Geltrex (Gibco)-coated wells. Routine cell passages were carried out using ReLeSR (Stem Cell Technologies) to detach cells. For experiments, cells were dissociated into a single-cell suspension using Accutase (Gibco).

*Establishing synNotch-KOLF2.1J iPSC line*: KOLF2.1J iPSCs were engineered to stably express synNotch platform elements (i.e., receptor and transgene payloads) via a Sleeping Beauty transposase/transposon system. Using the TransIT®-LT1 Transfection Reagent (Mirus, #MIR 2300), 1 µg of the Sleeping Beauty 100x transposase (Addgene 34879, a kind gift from Zsuzsanna Izsvak) 46 and 1 µg of the transposon containing the synNotch receptor and transgene payloads were transfected into the iPSCs. Approximately 1 x $10^6$ cells / mL were plated into a Geltrex-coated well in a 6-well plate and allowed to incubate with the DNA complexes for 24 hours at 37°C in StemFlex medium supplemented with 10 µM Y-27632 ROCK inhibitor (Tocris, #1254). After 24 hours, medium was replaced with fresh media. Engineered cells were selected with 0.6 µg/mL puromycin during sub-cultivation and sorted at the Vanderbilt Flow Cytometry Shared Resource with a 4-laser FACSAria III based on receptor expression using a c-myc-tag epitope appended to the synNotch receptor.

*Activation of synNotch engineered iPSCs*: To activate the GFP-responsive, LaG16 synNotch iPSCs, we supplemented StemFlex media with 0, 1, 5, 10, 50, 100, 500, or 1000 nM of dimeric EGFP. Dimeric EGFP was supplemented in the media when cells were plated and mCherry expression was assessed after 48 hours. To quantify mCherry expression, cells were fixed with 4% paraformaldehyde in DPBS for 10 minutes, blocked and (5% FBS and 0.3% Triton X in DPBS), and incubated with mCherry primary antibody (Supplemental Table 2) overnight at 4ºC. Cells were then washed three times with DPBS before incubation with an Alexa Fluor 647-conjugated Donkey anti-rabbit IgG secondary antibody (Supplemental Table 2) and the nuclear stain DAPI (Thermo Scientific, 62247, 1:1000). Cells were protected from light and incubated for 1 hour at room temperature. Cells were then washed with DPBS three times prior to imaging cells on a Leica Dmi8 microscope.

*ImageJ analysis*: Mean mCherry pixel intensity was calculated using ImageJ/Fiji software (available from the NIH at https://imagej.net/software/fiji/downloads) using built-in functions. Briefly, fluorescent images were converted to 8-bit greyscale images. DAPI channel images were used to develop a region of interest (ROI) that was used to measure the mean grey value in the mCherry channel. The ROI was made by subtracting out background noise (rolling ball radius = 50) and thresholding the greyscale image (85,255) to make a binary mask. Nuclei were then defined using the watershed feature prior to using the default analyze particles function to add the DAPI mask to the ROI manager. The ROI was then overlayed on the greyscale mCherry images to calculate the mean grey value for each nucleus. These values were then averaged to obtain the mean mCherry pixel intensity per nucleus.

**Supplementary Table 1.** Antibody variable fragments used to produce synNotch receptors.

| scFv | Target | $K_d$ |
|---|---|---|
| TNF3E[10] | TNF | 16pM |
| PET1074B9[11] | TGF-β1 | Not reported |
| G6-31[12,13] | VEGF | 0.9nM |
| Canakinumab[14] | IL-1β | 4.1nM |
| Ziltivekimab[15] | IL-6 | 0.52nM |
| Gevokizumab[14] | IL-1β | 0.29nM |
| Olokizumab[16] | IL-6 | 10pM |

**Supplementary Table 2.** Antibodies used for immunolabeling.

| Antibody Target-label | Vendor | Dilution | Catalog # |
|---|---|---|---|
| Myc-Alexa Fluor 647 | Cell Signaling Technology | 1:50 | 2233S |
| Mouse CD86-PE | Biolegend | 1:800 | 159203 |
| Mouse CD163-Brilliant Violet 421 | Biolegend | 1:400 | 155309 |
| Mouse CD206-Pacific Blue | Biolegend | 1:400 | 141707 |
| mCherry-Rabbit IgG | Cell Signaling Technology | 1:200 | 43590 |
| Donkey anti-Rabbit IgG-Alexa Fluor™ 647 | Thermo Fisher Scientific | 1:500 | A31573 |

**Supplemental Figures**

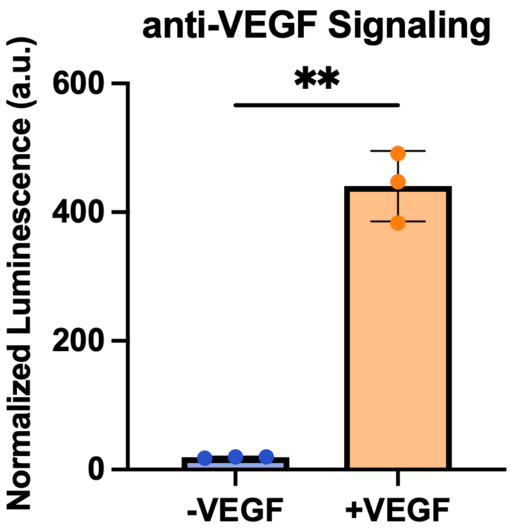


**Supplementary Figure 1.** Material independent VEGF synNotch signaling. L929 fibroblasts engineered with anti-VEGF synNotch demonstrate ligand dependent activation when supplemented with 25ng/mL of VEGF. Firefly luminescence readout normalized to cell viability. A Welch's t test of significance was performed. n=3 for all experimental groups $^{**}p<0.01$.

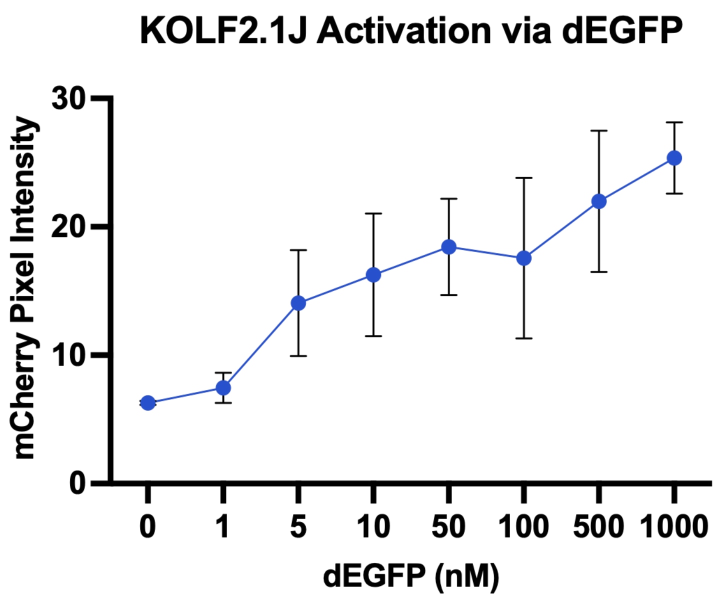


**Supplementary Figure 2.** KOLF2.1J iPSC dimeric EGFP (dEGFP) activation. dEGFP dose response in KOLF 2.1J human induced pluripotent stem cells (iPSCs) engineered with a GFP-responsive LaG16 synNotch receptor and an inducible mCherry fluorescent protein payload. Cells were activated with 0-1000nM of dEGFP for 48 hours before being fixed and immunolabeled for mCherry expression and the nuclear marker DAPI. Cells were imaged and mean mCherry pixel intensity was quantified using ImageJ software. Error bars indicate SEM. *n* = 15, 5 images from 3 individual wells.

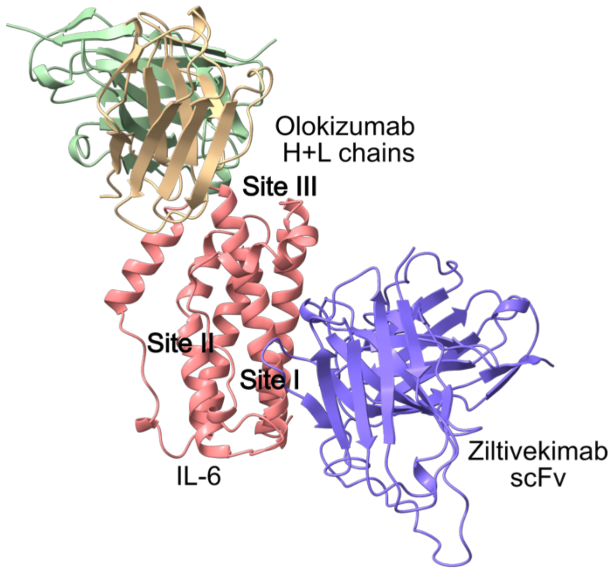


**Supplementary Figure 3.** Alphafold Multimer model of dual epitope binding of olokizumab and ziltivekimab variable chains to IL-6. AlphaFold multimer[1–7] modeling of IL-6 (Red) with ziltivekimab scFv (Blue)and olokizumab heavy (Green) and light (yellow) chains. Graphic demonstrates predicted ziltivekimab binding to site I of IL-6 as hypothesized and olokizumab binding to site III of IL-6 as described previously.[8,9] Model quality scores: pLDDT=91.7 pTM=0.844 ipTM=0.808

### Supplementary Figure 4: Figure 4 synNotch and SNIPR Flow Sort Plots

#### Wild-type mMSC

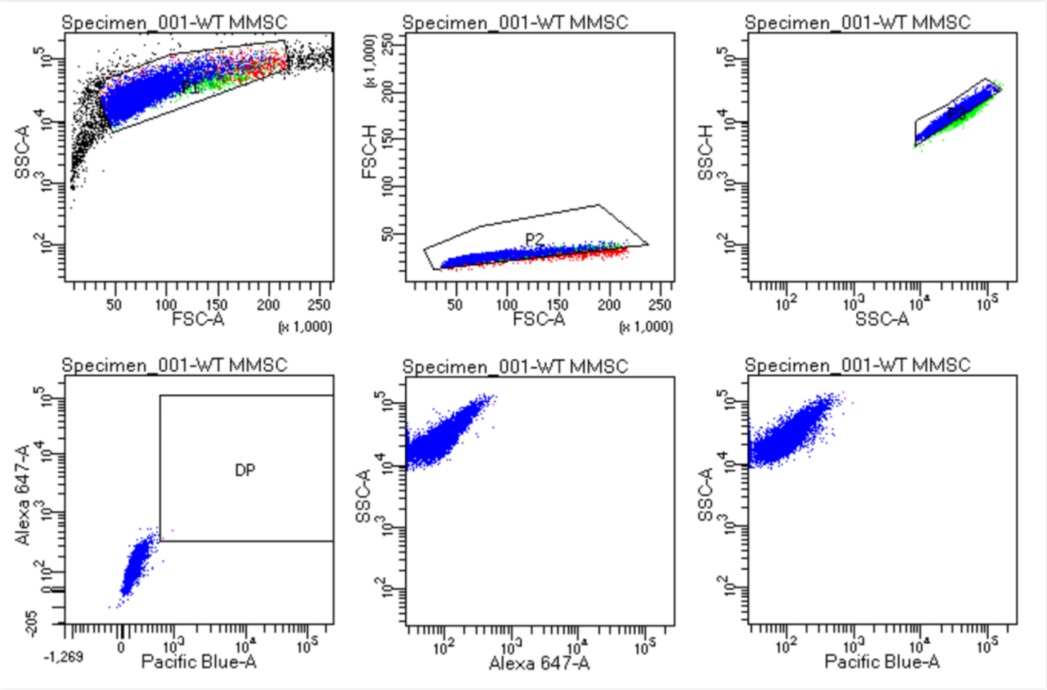


#### Canak-Notch mMSC

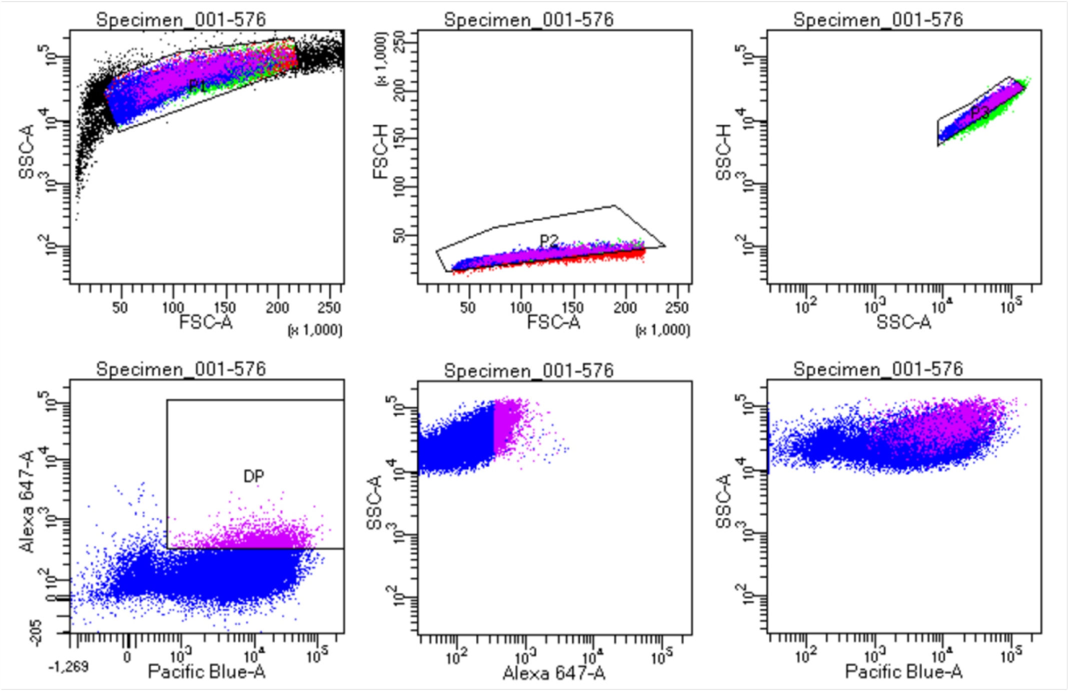

**Canak-SNIPR mMSC**

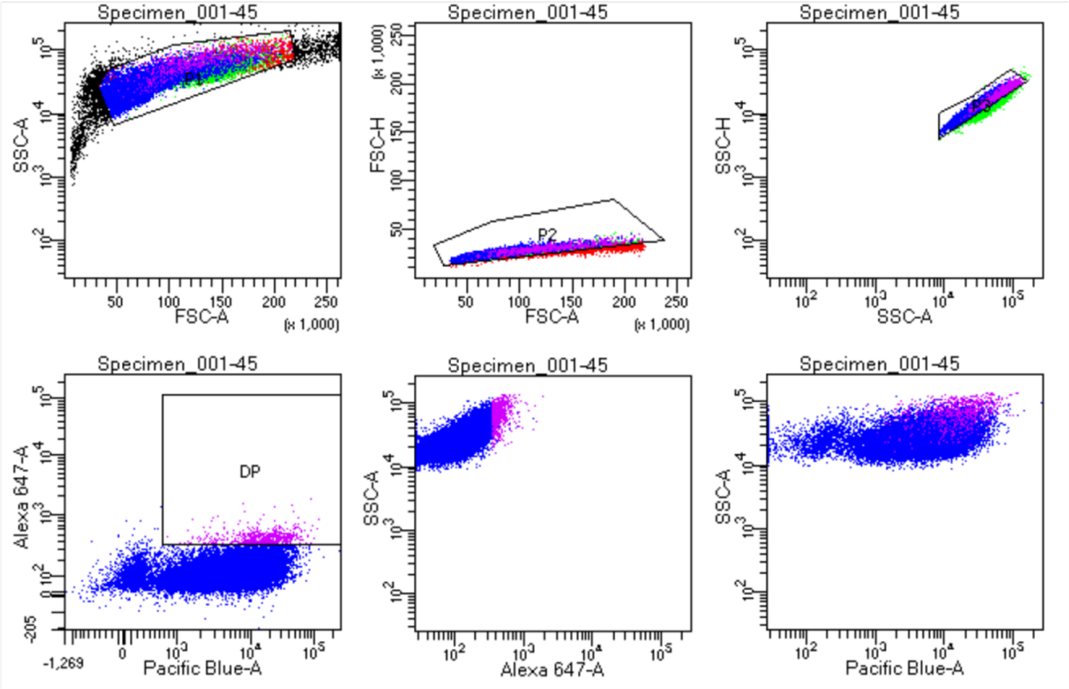


**Zilti-Notch mMSC**

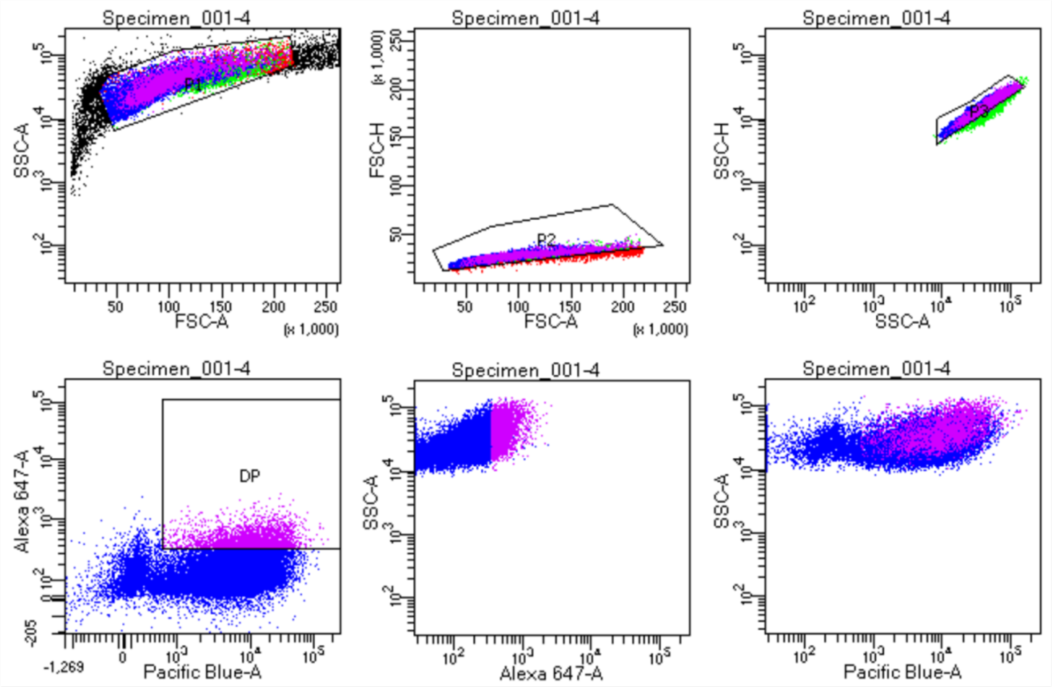

**Zilti-SNIPR mMSC**

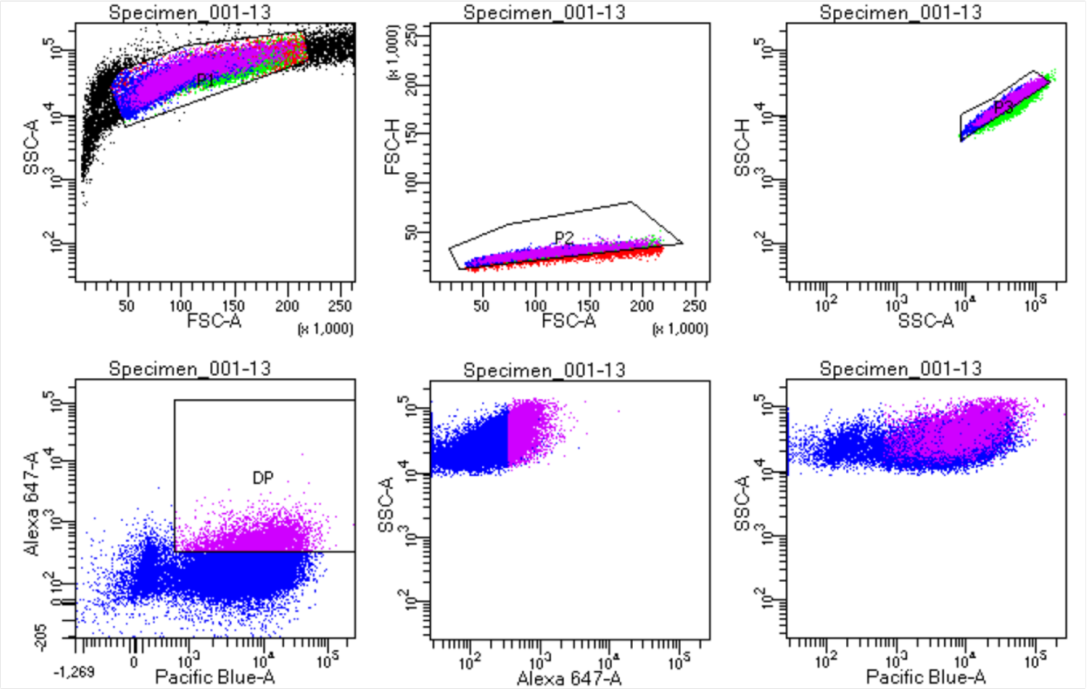


**Supplementary Figure 4.** Fluorescence activated cell sorting plots and gates for Figure 4. Alexa 647 indicates receptor expression detected via immunolabeling of an N-terminal c-myc epitope tag, and Pacific Blue indicates payload cassette expression via constitutive tagBFP expression. Plots show similar levels of expression between receptor constructs.

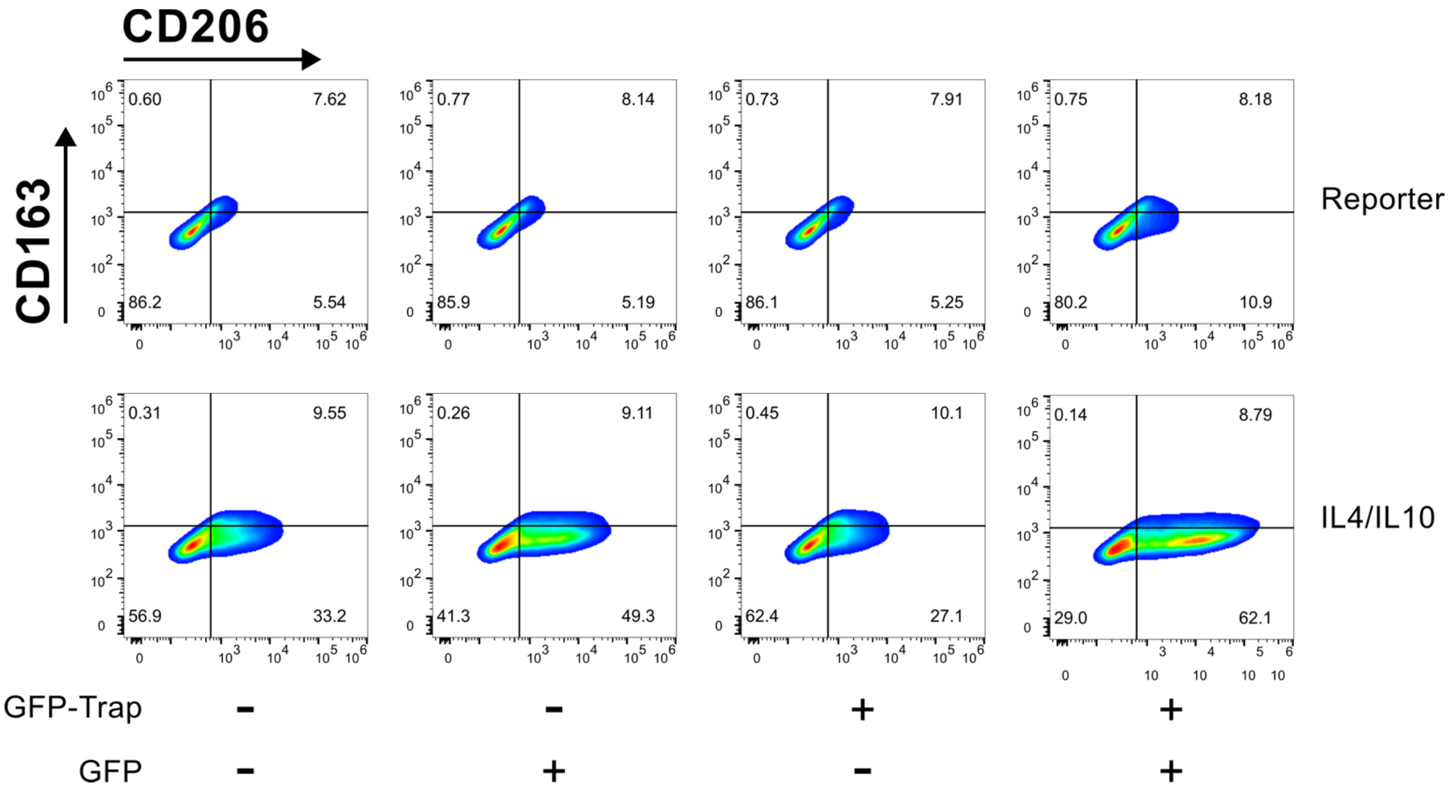


**Supplemental Figure 5.** GFP mediated RAW264.7 polarization flow plots**.** CD206 (x-axis) and CD163(y-axis) flow plots of GFP-mediated polarization experiment.

**Supplementary Figure 6: IL-1β mediated RAW264.7 polarization flow plots**

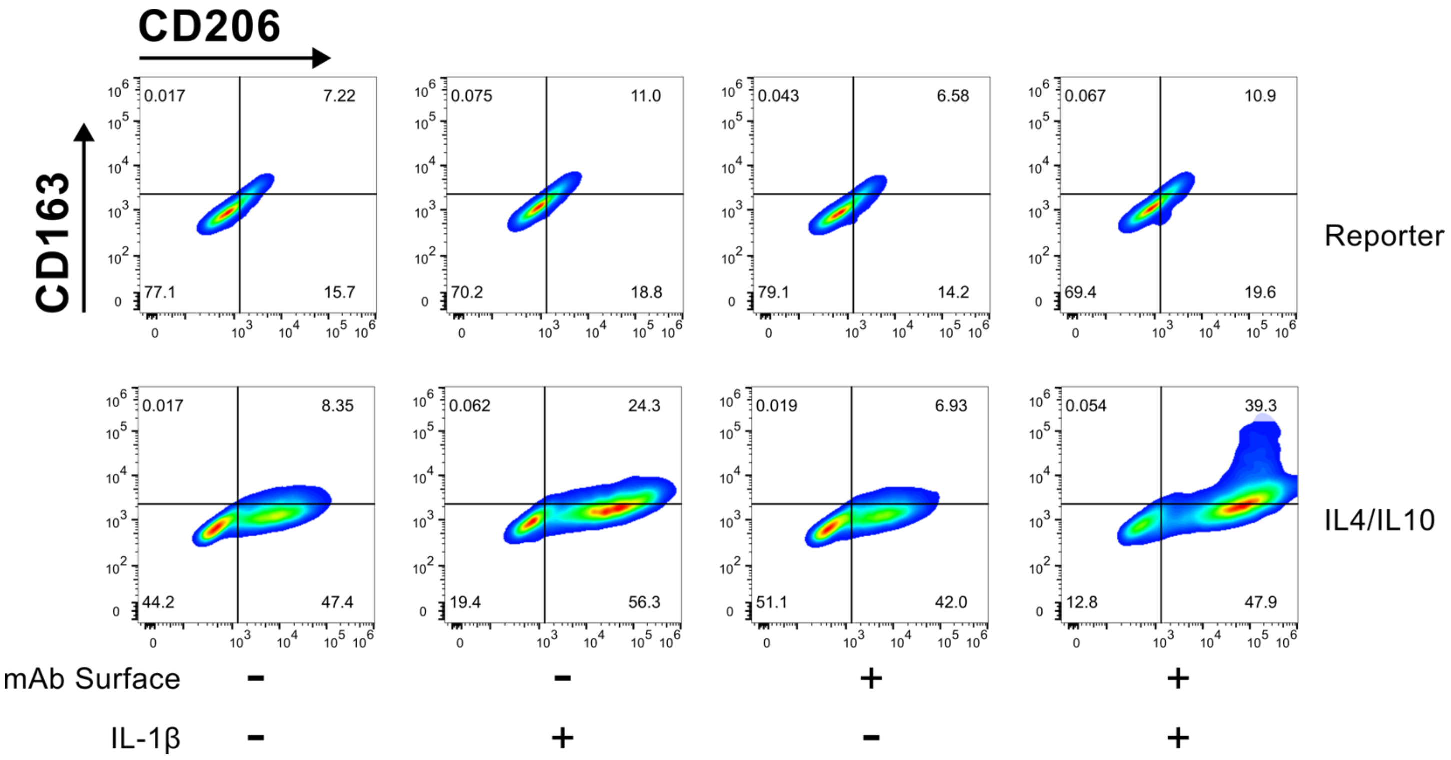


**Supplemental Figure 6.** CD206 (x-axis) and CD163(y-axis) flow plots of IL-1β-mediated polarization experiment.
